# Why did the animal turn? Time-varying step selection analysis for inference between observed turning points in high frequency data

**DOI:** 10.1101/2020.08.13.249151

**Authors:** Rhys Munden, Luca Börger, Rory P. Wilson, James Redcliffe, Rowan Brown, Mathieu Garel, Jonathan R. Potts

## Abstract

1. Step selection analysis (SSA) is a fundamental technique for uncovering the drivers of animal movement decisions. Its typical use has been to view an animal as “selecting” each measured location, given its current (and possibly previous) locations. Although an animal is unlikely to make decisions precisely at the times its locations are measured, if data are gathered at a relatively low frequency (every few minutes or hours) this is often the best that can be done. Nowadays, though, tracking data is increasingly gathered at very high frequencies, often ≥1Hz, so it may be possible to exploit these data to perform more behaviourally-meaningful step selection analysis.
2. Here, we present a technique to do this. We first use an existing algorithm to determine the turning-points in an animal’s movement path. We define a “step” to be a straight-line movement between successive turning-points. We then construct a generalised version of integrated SSA (iSSA), called *time-varying iSSA* (tiSSA), which deals with the fact that turning-points are usually irregularly spaced in time. We demonstrate the efficacy of tiSSA by application to data on both simulated animals and free-ranging goats (*Capra aegagrus hircus*), comparing our results to those of regular iSSA with locations that are separated by a constant time-interval.
3. Using (regular) iSSA with constant time-steps can give results that are misleading compared to using tiSSA with the actual turns made by the animals. Furthermore, tiSSA can be used to infer covariates that are dependent on the time between turns, which is not possible with regular iSSA. As an example, we show that our study animals tend to spend less time between successive turns when the ground is rockier and/or the temperature is hotter.
4. By constructing a step selection technique that works between observed turning-points of animals, we enable step selection to be used on high-frequency movement data, which are becoming increasingly prevalent in modern biologging studies. Furthermore, since turning-points can be viewed as decisions, our method places step selection analysis on a more behaviourally-meaningful footing compared to previous techniques.

## 1 Introduction

Understanding the factors that drive animal movement is a cornerstone of movement ecology (Nathan et al., 2008). Step selection analysis (SSA) is a powerful and well-used tool for providing such understanding, by identifying the habitat selection decisions that shape an animal’s movement patterns (Fortin et al., 2005; Rhodes et al., 2005). Recently, integrated Step Selection Analysis (iSSA) (Avgar et al., 2016) was introduced to generalise SSA. This provides a solid mathematical foundation for step selection analysis studies and enables inference of how landscape features influence both the movement and habitat selection of animals.

So far, the principal use of SSA (and iSSA) has been on data gathered at a sufficiently low frequency that it is reasonable to suppose an animal makes a distinct choice to move between successive location measurements, with GPS data being perhaps the prime example. This is despite the fact that, in reality, this remains a rather strong and often biologically unrealistic assumption, as decisions to turn are not regularly spaced in time. Recently, however, tracking technology has begun to allow scientists to gather data at sufficiently high frequencies that the resulting data is essentially continuous, since the distance between successive location fixes of an animal is often less than its body length. Data from magnetometers and accelerometers have allowed path reconstruction at sub-second frequencies, often over long periods of time, such as weeks or months (Wilson et al., 2008, 2013a; Street et al., 2018). Alongside this, accelero-magnetometer data can be used to understand energy expenditure (Wilson et al., 2020), classify behaviours (Yoda et al., 1999; Moreau et al., 2009; Nathan et al., 2012), and gain insight into an animal’s internal state (Wilson et al., 2014; Downey et al., 2017; Kröschel et al., 2017). These insights have the potential to be combined with location data to uncover details of what drives animal movement decisions in greater detail than ever before (Williams et al., 2020).

In comparison, GPS data, typically gathered at much lower resolutions, may not give a good indication of an animal’s behavioural decisions over short temporal scales (Hebblewhite and Haydon, 2010). Although some GPS data can now reach high frequencies (1Hz) (Ryan et al., 2004), the battery lifespan is greatly reduced, since GPS-based systems are power hungry. This leads to either a decrease in the duration of the study or an increase in battery size and therefore tag weight, which is limited to be a relatively small proportion of the animal’s body weight so as not to have too great an effect on the animal’s behaviour (Rasiulis et al., 2014). (This proportion is often cited as < 3% (Kenward, 2000), but can be higher for some animals (Barron et al., 2010).) Transmission telemetry is limited by the environment whereas biologging systems (such as accelero-magnetometers) are not, which means that biologgers can be applied across a wide range of taxa from marine (Tanaka et al., 2001; Noda et al., 2014) to aerial (Shepard et al., 2008; Williams et al., 2015) to terrestrial (Bidder et al., 2012; Street et al., 2018).

However, it is not always simple to adapt existing statistical techniques for use with high-frequency biologging data, since such techniques were often developed with lower frequency data in mind. For example, it does not make sense to apply SSA to ‘steps’ between successive measurements which may be less than a second apart, as is often the case with accelero-magnetometer data, since such steps cannot reasonably be viewed as representing distinct behavioural decisions of an animal. Instead, over the majority of such ‘steps’, an animal will most likely just continue to follow an already chosen path.

This issue of scale disparity between the actual decisions made by animals and the data gathered on them has a long, and occasionally controversial, history. An early study to address this was that of Turchin et al. (1991), who stressed the importance of demarcating the actual turns made by an animal when analysing movement paths (see also Turchin (1998)). Several further studies have shown that failure to account properly for scale in animal movement can lead (and has sometimes led) to misinterpretation of the properties of movement paths (Benhamou, 2014; Nams, 2005; Plank et al., 2013; Turchin, 1996). Perhaps a partial reason for this is that locational data is often not sufficiently high frequency for users to be able to analyse a wide variety of scales, especially at the level of the fine-grained ‘moves- and-turns’ analysis originally studied by Turchin et al. (1991). However, modern biologging offers the opportunity to begin resolving these issues by demarcating movement decisions from high-resolution paths.

As a first stage towards this end, Potts et al. (2018) proposed an algorithm (referred to here as the Turning-Points Algorithm) which identifies the key turning-points in an animal’s path. Using this, we can consider the movement from one turning-point to the next as a ‘step’. (Note that the movement between successive turning-points was called a ‘move’ by Turchin (1998), but we will use the term ‘step’ to correspond with the phrase ‘step selection analysis’.) These steps likely correspond to actual movement decisions made by the animal: since turns are energetically costly (Wilson et al., 2013b), they would only be made if there were a benefit sufficient to outweigh the costs of turning. However, unlike applications of SSA to successive, regularly-gathered GPS locations, steps between turning-points are not evenly spaced in time.

In principle, it is possible to ignore time and just use the SSA (or iSSA) method on the steps between successive turning-points. However, there are disadvantages to this approach. One is that certain movement covariates may depend upon the time between successive turns, a quantity we will call the *step-time*, by analogy to the related concept of step-length. For example, animals may tend to have a longer time between turns in open environments compared to those that are more difficult to cross. Failure to incorporate the effect of covariates on step-times could potentially lead to inaccurate inference of behaviour. In addition, the resulting movement kernel would not describe a model that explicitly incorporates time, so it could not be propagated forwards in time to make predictions of long-term space-use patterns (e.g. home ranges; Börger et al. (2008)). Yet prediction of broad-scale space use is one of the key aims of many recent studies in step selection analysis (Potts et al., 2014a,b; Avgar et al., 2016; Merkle et al., 2017; Signer et al., 2017; Potts and Schlägel, 2020). Ideally, the SSA procedure should be adapted to accommodate explicitly for the non-constant step-times inherent in paths described by the actual turns of the animals. The principle technical advance of this work is to enhance iSSA so it can be used on data with such non-constant step-times. We call this method time-varying iSSA (tiSSA).

To demonstrate the efficacy of tiSSA, we apply it to 1Hz dead-reckoned data (1 location per second) from a group of free-ranging goats (*Capra aegagrus hircus*) during the summer and living in an alpine pasture surrounded by steep slopes in the French Alps. They spend a large part of their time (roughly 40% of the study duration) in a relatively small region (radius approximately 70m) around a central area, which encompasses a goat pen and nearby salt-licks, from which they make regular foraging excursions to graze and browse along the mountain slopes. As a basic demonstration of our method, we make three simple hypotheses about goat movement: that they will have a tendency to (i) display a preference to move toward the central area, where the strength of attraction is greater the further they are away from it, (ii) choose smaller-angled turns than larger (i.e. have persistent movement), and (iii) avoid steep-sided terrain, since this requires more energy expenditure to traverse (Minetti et al., 2002; Ardigò et al., 2003). Whilst these hypotheses are intended principally as a proof of concept for the tiSSA algorithm, it is worth noting that the third hypothesis has a plausible alternative: that goats may indeed prefer steep sided terrain to reach areas of high elevation where they can take refuge from predators. Our analysis for (iii) can thus be viewed as testing between these competing hypotheses.

In addition to these habitat-and movement-selection covariates, we also demonstrate the use of tiSSA for inferring environmental drivers of the duration of step-times. Visual observations suggest that the goats have more directed paths when moving back toward the central area. Furthermore, when the temperature is high, they tend to restrict their area of movement to the shaded regions near the pen and other nearby buildings. We also noticed the goats seem to find it more difficult to move in rocky terrain, so make more turns. Inspired by these observations, we give a simple demonstration of the utility of incorporating time-dependent covariates with three further hypotheses: (a) that step-times are longer when the goats are moving toward the central point, (b) that step-times are shorter when the temperature is higher, and (c) that step-times are shorter when the goats are moving through rocky terrain.

We further demonstrate the value of tiSSA by comparing it with the traditional use of SSA (or iSSA) (Fortin et al., 2005; Rhodes et al., 2005; Avgar et al., 2016), whereby the step-time between successive location fixes is constant. For this, we use both the aforementioned data on goat movement and also simulated paths. We subsample our paths at a variety of constant step-times and compare the inference using these subsampled paths with that from the paths defined by the Turning-Points Algorithm. We investigate how the accuracy and precision of inference varies as the subsampling interval is changed, thus demonstrating how step selection with constant subsampling of a movement path (as is typical in many SSA or iSSA studies) may give rise to misleading results.

In summary, our study both (a) shows the great value in gathering high frequency data to understand the drivers of animal movement, and (b) gives a usable method for making behavioural inferences with such data, where the inference is now drawn from movements between behaviourally meaningful points: the points where the animal has actually made a decision to turn.

## 2 Methods

### 2.1 Data

Data on goat (*Capra aegagrus hircus*) movement were collected at the Bauges Mountain (Massif des Bauges, 45.61°N, 6.19°E) of the French Alps, using triaxial accelerometers and magnetometers (Wildbytes Technologies http://www.wildbyte-technologies.com) contained in Daily Diary tags (Wilson et al., 2008), combined with Gipsy5 GPS tags (TechnoSmArtTracking Systems http://www.technosmart.eu) inside custom-built 3D printed ABS plastic housings attached to commercial nylon collars (Fearing Lifestyles, Durham, UK). Accelerometer data were collected at a frequency of 20 Hz, magnetometer at 8 Hz, and temperature at 2 Hz. GPS locations were collected every 15 minutes. The data were dead-reckoned at 1 Hz (Bidder et al., 2015) with the Framework4 software (Walker et al., 2015) to reconstruct paths of locations over time. Overall, seven trajectories were reconstructed from the data, each of which was one week long.

The goats tended to spend most of their time in a central area (radius ~ 70m) which contains a pen and nearby salt licks. We define the centre of this area as the *central point* for the purposes of this manuscript. As well as locational data, we also use data on the terrain and elevation. The terrain consists of areas of scree biotope (Devillers et al., 1991), which we term *rocky terrain*, as well as woodland and grassland. Elevation recordings were found using Google’s Elevation API (https://developers.google.com/maps/documentation/javascript/elevation) at a resolution of 1m.

### 2.2 Time-varying integrated step selection analysis (tiSSA)

The central aim of this work is to demonstrate the benefit of using high frequency data for SSA. Usually, SSA is used to analyse the movement choices between successive recorded locations. However, for data collected at very high frequencies (e.g. ≥ 1Hz), the distances between successive locations are so short that we cannot expect to infer useful behavioural information by applying SSA to every possible step in the path. Instead, we use a pre-processing method to find those locations on each path where the animal appears to have made a distinct decision to turn, which are more interesting from a behavioural viewpoint than regularly (in time) sampled locations, before applying the SSA procedure.

Specifically, we use the method of Potts et al. (2018) to identify these turning points, thus simplifying an animal’s path into a series of turning-points and straight line segments joining these points (Fig. 1a-b). These straight line segments will then be the “steps” in our subsequent analysis. Each step has both a step-length and a step-time, where the latter is defined to be the time between successive turns. Different from the way SSA and iSSA have usually been used in previous work, the step-times will not all be the same. Thus we need to modify the iSSA method to incorporate this.

**Figure 1.**
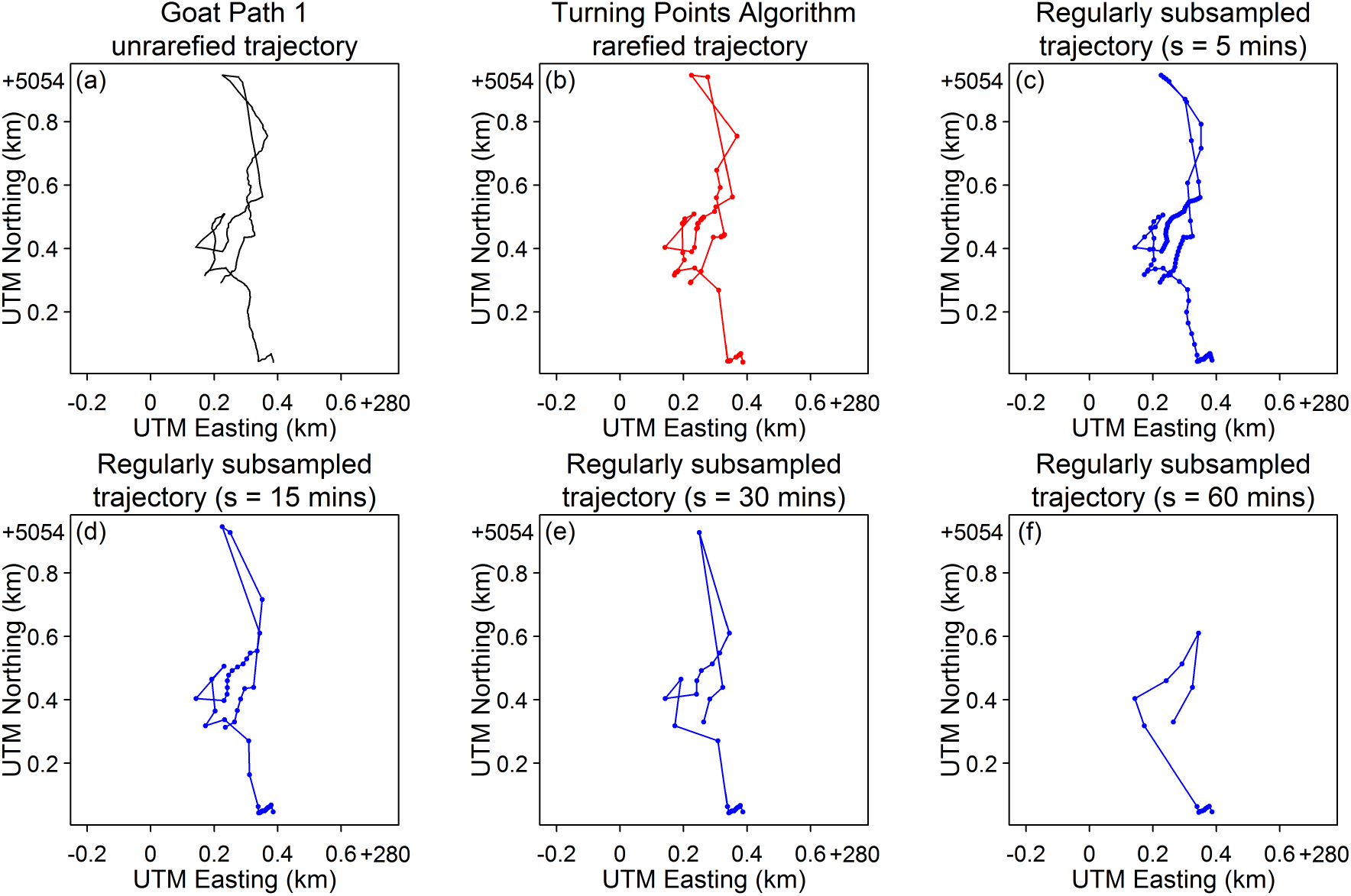
A comparison of a path constructed at a frequency of 1Hz to the same path rarefied (a) using the Turning-Points Algorithm and (b) by subsampling at regular time intervals (c-f).

The tiSSA method, which we now introduce, is designed to make this required extension to iSSA (Avgar et al., 2016). We start by defining the *movement kernel*, which is the probability density function (pdf) of the animal making its next turn at location **x** after a time *τ* has elapsed, given that it is currently making a turn at location **z** and arrived at **z** on a bearing of *α*_**z**_. This has the following functional form

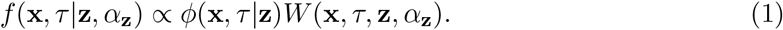

Here, ∝ means ‘proportional to’, and the associated constant of proportionality is defined so that *f* (**x**, *τ*|**z**, *α*_z_) integrates to 1 over all possible values for **x** and *τ* (i.e. *f* is a probability density function). The function *ϕ* is a kernel of available step-lengths and times, which we refer to here as the *sampling kernel*, whilst *W* is called the *weighting function*. Note that the sampling kernel here is equivalent to the function *g* in Equation (2) of Avgar et al. (2016) or *ϕ*\* in Forester et al. (2009). It should not be confused with the ‘resource-independent movement kernel’ from Forester et al. (2009) or the ‘movement kernel’ Φ from Avgar et al. (2016), which are different entities (indeed, later on we explain how to construct the resource-independent movement kernel for tiSSA).

In iSSA (or SSA) the step-times are constant, so the sampling kernel, *ϕ*, is a distribution of step-lengths and turning angles. Here, however, since the step-times are non-constant, we also need to incorporate a distribution of step-times into *ϕ*. We thus define the sampling kernel as

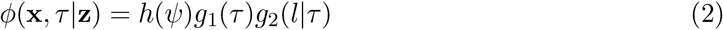

where *l* = |**x** − **z**| is the step-length, *ψ* = *H*(**x**, **z**) is the heading from **z** to **x**, *h*(*ψ*) is the distribution of headings, *g*_1_(*τ*) is the distribution of step-times, and *g_2_(l|τ*) is the distribution of step-lengths given a particular step-time (which is equivalent to specifying a distribution of step-speeds). Avgar et al. (2016) suggest using a gamma distribution for the step-lengths, due to its generality (e.g. the exponential and *χ*^2^ distributions are special cases). Similarly, we use gamma distributions for both *g*_1_(*τ*) and *g*_2_(*l*|*τ*).

The pdf of a gamma distribution can be written in exponential form as follows

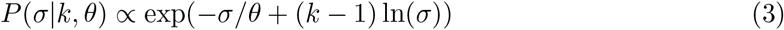

where *k* and *θ* are the shape and scale parameters, respectively, and *σ* represents either the step-time, *τ*, or step-length, *l*. This particular way of writing the gamma distribution is useful during the inference procedure, below. When performing step selection analysis with non-constant step-times, we sample randomly from the kernel given by Equation (2). This gives control locations for each start position, which can be compared with the measured case location using conditional logistic regression, following the procedure given in, for example, Avgar et al. (2016). Mathematical justification for using this conditional logistic regression procedure to parametrise Equation (1) is given in Supplementary Appendix S1.

The weighting function has the following form

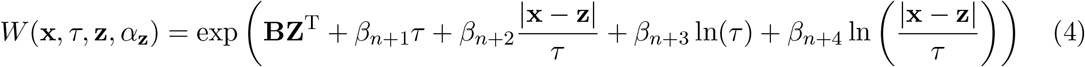

where **Z** = [*Z*_1_(**x**,**z**, *α*_z_,*τ*), *Z*_2_(**x**,**z**,*α*_z_,*τ*),…, *Z_n_*(**x**,**z**, *α*_z_,*τ*)] is a vector of covariates and **B** = [*β*_1_,*β*_2_,… *β_n_*] is a vector of coefficients representing the effect of each covariate on movement decisions (Avgar et al., 2016). Here, *β*_*n*+1_ and *β*_*n*+3_ correct for the step-time, whilst *β*_*n*+2_ and *β*_*n*+4_ correct for the step-speed. Notice now the reason for writing the gamma distribution in the form of Equation (3): *β*_*n*+1_ and *β*_*n*+2_ correct the scale parameter of the time and speed respectively, whilst *β*_*n*+3_ and *β*_*n*+4_ correct the shape parameter of the time and speed respectively. Including these correcting factors is important to avoid potentially biased results (Forester et al., 2009; Avgar et al., 2016). Furthermore, it is now possible to define the resource-independent movement kernel, *sensu* Forester et al. (2009), as the product of *ϕ*(**x**, *τ*|**z**) from Equation (1) and exp(*β*_*n*+1_*τ* + *β*_*n*+2_|**x** − **z**|/*τ* + *β*_*n*+3_ ln(*τ*) + *β*_*n*+4_ln(|**x** − **z**|/*τ*)) from Equation (4).

The covariates, *Z_i_*, may depend on the end of the step, **x**, the start of the step, **z**, along the step (between **z** and **x**), the direction the animal is moving in when it arrives at **z** and/or the time it takes to move from **z** to **x**, given by *τ*. Consequently the *Z_i_* are functions of **x**, **z**, *α_z_* and *τ*. Each *Z_i_*(**x**, **z**,*α*_z_,*τ*) represents a statement about what drives the animal’s decision to move.

### 2.3 Application to empirical data

We use tiSSA on paths reconstructed from the goat movement data using the Turning-Points Algorithm (Potts et al., 2018). To compare our method with the traditional use of iSSA, which uses data sampled regularly at a relatively low resolution (e.g. every few minutes or hours), we subsample each of our paths at regular time-intervals, referred to as regular subsampling. For this, we use step-times (i.e. subsampling resolutions) ranging from 5 seconds to 420 minutes. Each of these step-times leads to a slightly different rarefied path. By considering the *β_i_*-values inferred using the paths from the Turning-Points Algorithm as indicative of the ‘real’ movement tendencies (denoted 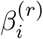), we assess the accuracy of the *β_i_*-values inferred using regular subsampling (denoted 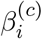), by comparing each 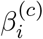 to the corresponding 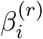.

#### 2.3.1 Non-constant step-times

To apply tiSSA to the paths inferred using the Turning-Points algorithm, we use a uniform distribution for the headings and gamma distributions for the step-times and step-lengths. In other words, we define

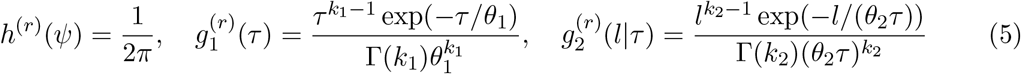

where *k*_1_ and *θ*_1_ are, respectively, the shape and scale parameters of the gamma distribution fitted to the step-times, and *k*_2_ and *θ*_2_ are the best fit shape and scale parameters (respectively) for the probability density function 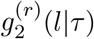. (We also show that the same *k*_2_ and *θ*_2_ are the best fit shape and scale parameters for a gamma distribution of step-speeds in Supplementary Appendix S2, which can ease inference of these parameters). We use the superscript (r) to represent functions and coefficients used for non-constant step-time data.

Substituting Equation (5) into Equation (2) gives the following

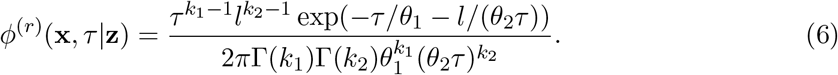

To build our weighting function, *W*^(*r*)^(**x**, *τ*, **z**, *α*_z_), we test various hypotheses, namely that goats tend to

(A1) move toward a central point (**x**_cp_) with strength proportional to their distance from this central point,
(A2) have longer step-times when both further away from the central point and moving towards it, since goats returning should be minimising deviations,
(B) persist in the same direction,
(C) avoid steep upward trajectories,
(D) have shorter step-times when the temperature is higher, since goats in hot weather should be exhibiting movement that keeps them in the shade,
(E) have shorter step-times when moving through rocky terrain, since goats moving over topographically challenging substrate should be selective in their tracks which leads to them turning frequently.

Notice that A1 and A2 should not be tested within the same weighting function, as they conflate with one another (hence the labelling). Each of these hypotheses has a corresponding covariate. These covariates have the following functional forms

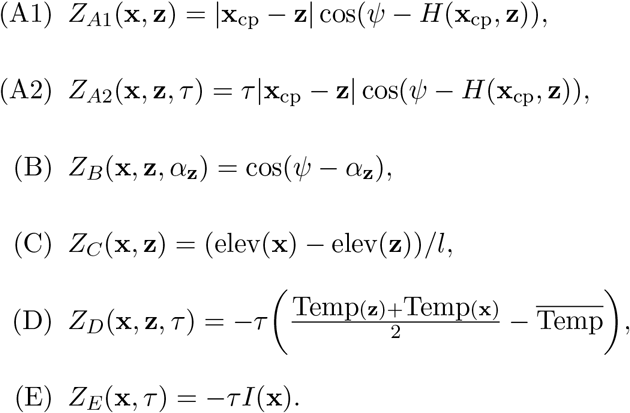

Here, elev(**x**) is the elevation at **x** (found using Google’s Elevation API), Temp(**x**) is the temperature recorded from the collar when the goat was at **x**, 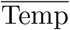 is the average temperature across all recordings, *I*(**x**) is an indicator function equaling 1 when **x** is in rocky terrain (as defined in Section 2.1) and 0 otherwise, and **x**_cp_ is the location of the central point. We note that there maybe a non-linear relationship between temperature and step-time, but here we assume that the relationship is linear for simplicity.

The point **x**_cp_ is defined to be the centre of the single *site of interest* found using the method of Munden et al. (2019) for each path, where a site of interest is an area where an animal spends a disproportionately long time. Here, the single site of interest encompasses part of a goat pen and the area outside this goat pen where salt-licks are placed. More information on identifying the sites of interest can be found in Supplementary Appendix S3.

We create four separate models using different combinations of the above covariates (*Z*_*A*1_, *Z*_*A*2_, *Z_B_*, *Z_C_*, *Z_D_*, *Z_E_*) to form the weighting function under each model, together with the correcting factors for the step-time and step-speed. The first model, called the *Base Model*, consists of covariates *Z*_*A*1_, *Z_B_* and *Z_C_*. Since it does not include any step-time dependent covariates we use it to compare the results from using constant and non-constant step-time data. We adapt the Base Model to create the three other models. The *Straight Returns Model* is constructed by replacing the covariate *Z*_*A*1_ with *Z*_*A*2_. The *Temperature Model* consists of the Base Model and covariate *Z_D_*. The *Rocky Terrain Model* consists of the Base Model and *Z_E_*. These three latter models are used as examples to demonstrate the range of step-time related questions we are able to answer using ultra-high-resolution data rarefied with a biologically-relevant criterion given by the Turning-Points Algorithm.

To give an example functional form, under the Base Model the movement kernel is defined as

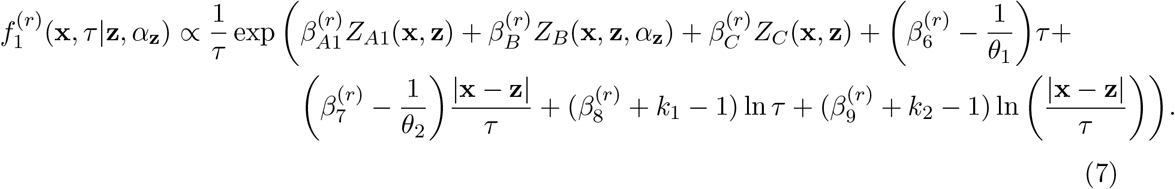

The functional forms of the movement kernels under the other models are given in Supplementary Appendix S4.

#### 2.3.2 Constant step-times

To perform iSSA on the regularly subsampled paths, and thus compare it with tiSSA plus the Turning-Points Algorithm, we define the sampling kernel as follows (Avgar et al., 2016)

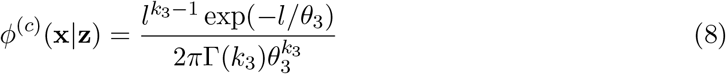

where *k*_3_ and *θ*_3_ are the shape and scale parameters from the gamma distribution fitted to the step-lengths. Note the difference between Equations (6) and (8): in the latter, no step-time distribution is required. The superscript (*c*) refers to any function or coefficient used with constant step-time data.

We use the same covariates as the Base Model above, but instead of correcting for the step-time 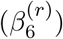 and step-speed 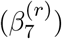 and the natural logarithm of each 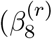 and 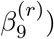, as we do for non-constant step-time data, we instead only correct for the step-length 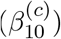 and its natural logarithm 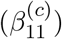. This leads to the following weighting function

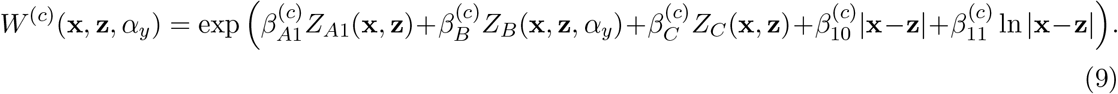

Note that in some paths and for certain step-times (namely path 4 for step-times of less than 5 minutes; paths 6 and 7 for step-times of less than one minute), there are steps of length zero, so in these cases we cannot correct for the natural logarithm of the step-length. In these cases, we omit 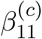.

The movement kernel is then

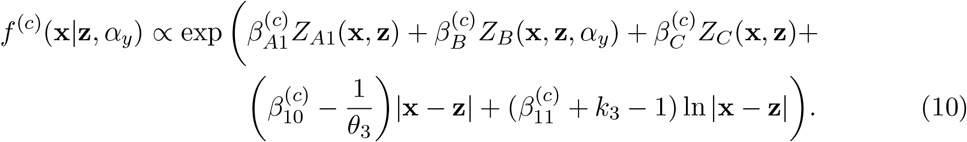

In all of our step selection analyses, both for constant and non-constant step-time data, we drew 100 control steps from the sampling kernel (*ϕ*^(*r*)^ or *ϕ*^(*c*)^ for non-constant or constant step-times, respectively) for each case step. For the non-constant step-time paths, drawing from the sampling kernel, *ϕ*^(*r*)^, is a two-step procedure, involving first drawing a step-time before a step-length. We use conditional logistic regression to find the *β_i_*-values, using the `clogit` function from the `R` package, `survival` (Therneau and Grambsch, 2000).

### 2.4 Application to simulated data

To test further the efficacy of tiSSA, as compared to iSSA using regularly subsampled locations, we constructed paths of simulated animals through a heterogenous environment. This is a valuable test as we have complete control over the individuals’ ‘movement decisions’, so we can compare inferred parameter values to the actual values used to construct the simulations. We cannot do this with empirical data, as we do not know the actual decisions made by real animals as they move.

To this end, we constructed nine paths, each consisting of 1,000 steps of an animal moving through the artificial resource layer given in Fig. 2a. Each step is constructed by drawing a new location **z**, given the current location **x**, from the following distribution

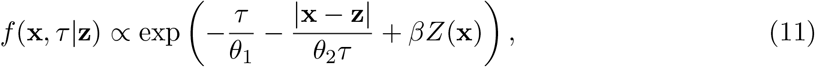

where *βZ*(**x**) is the value of the resource layer (Fig. 2a) at location **x**. Each of the paths consists of a different set of values for (*θ*_1_,*θ*_2_,*β*). We kept *θ*_1_ = 1 fixed, and used *θ*_2_ = 0.1,0.2,0.5, *β* = 0.5, 1, 2, leading to the nine aforementioned paths.

**Figure 2.**
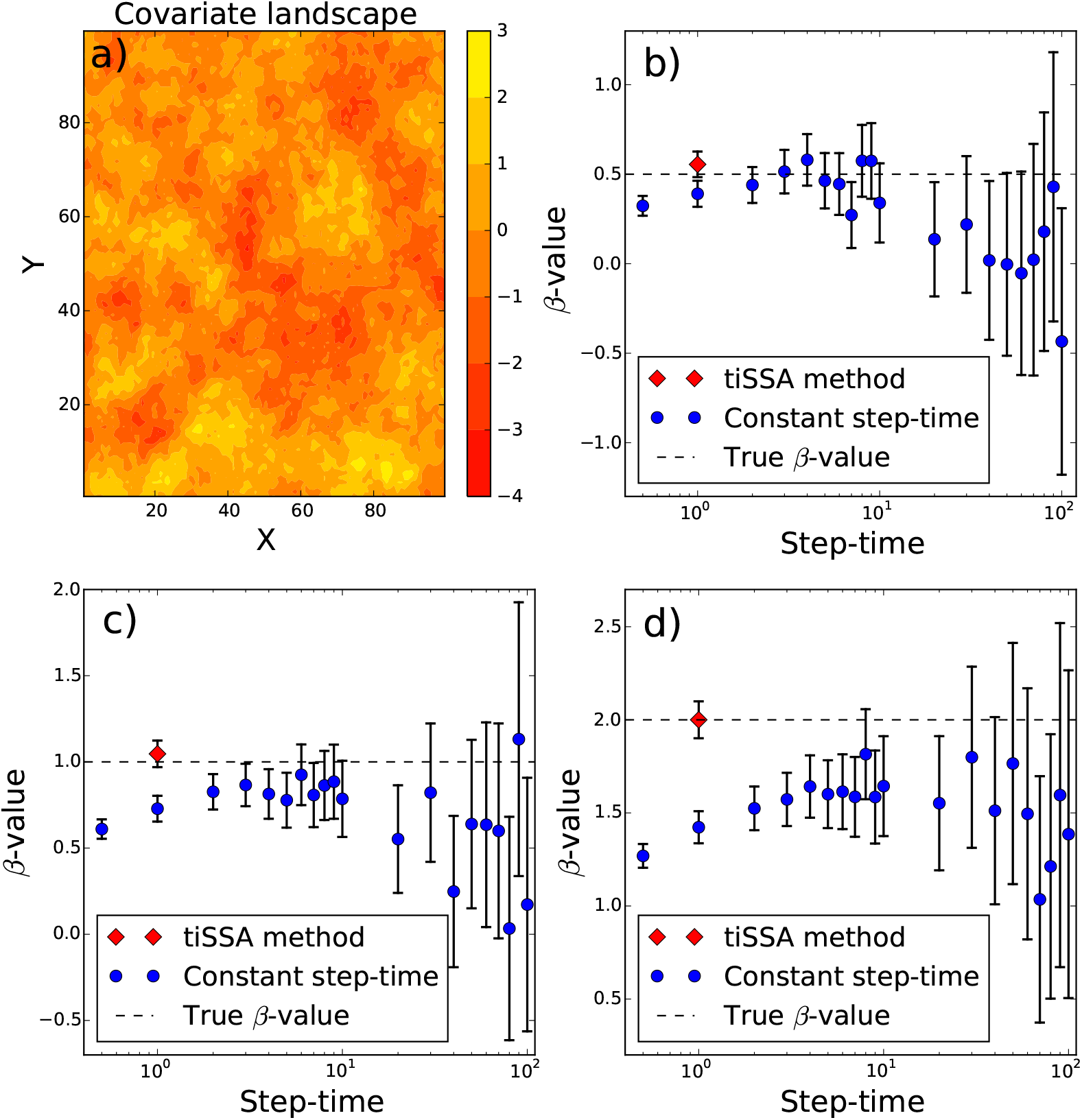
Results of the simulation analysis in the case *θ*_1_ = 1, *θ*_2_ = 0.1 (see Equation 11). Panel (a) shows the resource layer used to construct the simulated paths. Panels (b-d) show, respectively, the results for *β* = 0.5, *β* = 1, and *β* = 2. The red diamonds show results from analysing the paths using tiSSA, whereas the blue dots show results from using iSSA with paths subsampled at regular step-times. Error bars show 95% confidence intervals.

Having simulated the paths, we ran the tiSSA method over the set of points where the simulated animal had actually turned, to test whether tiSSA would return the true parameter values for (*θ*_1_,*θ*_2_,*β*). We also analysed the same paths using tiSSA but without including the covariates that correct for step-length and step-time (*β*_*n*+1_, …*β*_*n*+4_ from Equation 4), to test the importance of including these correction variables. We then subsampled each path at regular time intervals and used iSSA to infer the value of *β*, comparing this both with the results from tiSSA (similar to our analysis of the goat data) and the actual *β*-values used to construct the paths.

## 3 Results

For the simulated paths, the tiSSA analysis returned the correct *β*-value (within 95% confidence intervals) for all paths analysed (Fig. 2b-d, Supplementary Figure SF8). In contrast, the *β*-values returned by iSSA on regularly subsampled paths were often significantly different from the true *β*-values. Failure to correct for the step-length and step-time also led to incorrect inference of *β*. For example, when *θ*_2_ = 0.1 as in Fig. 2b-d, the 95% confidence intervals were (0.51, 0.58), (1.03, 1.11), (2.06, 2.15) for *β* = 0.5, 1, 2 respectively, when the corrections for step-length and step-time were not included. By correcting for these, however, the 95% confidence intervals were (0.48, 0.56), (0.97, 1.05), (1.90, 2.10) for *β* = 0.5, 1, 2 respectively. Full results are given in Supplementary Table ST2.

When applying tiSSA to the empirical paths on goat movements, we found that the tendency for avoiding steep upward trajectories is insignificant, both for the constant step-time paths and the paths created using the Turning-Points Algorithm, which may be due to the fact that the goats sometimes use areas of high altitude as refuges. Thus we removed the corresponding covariate (*Z_C_*) from all of our weighting functions in our subsequent analysis.

Fig. 3 shows results of analysing the Base Model with iSSA applied to constant subsampling (blue dots), compared with tiSSA applied to the paths rarefied by the Turning-Points Algorithm (red diamond; red dotted line), for an example path (see Supplementary Figures SF2-7 for the other paths). For *β*_*A*1_, which represents a tendency to move toward the central point, Fig. 3a shows that the inference from iSSA is highly dependent upon the frequency of the subsampling. The accuracy of inference (i.e. how close each blue dot is to the red dot in Fig. 3a) decreases as the frequency of subsampling moves away from the frequency of turns (as inferred by the Turning-Points Algorithm). However, even when the frequencies are approximately the same, the inferred *β*_*A*1_-value from constant subsampling (9.0 × 10^-4^ for a step-time of 10 minutes) is nearly twice that of the tiSSA method (4.9 × 10^-4^). When the step-time is 1 hour, the inferred *β*_*A*1_-value is more than 6 times higher (3.0 × 10^-3^).

**Figure 3.**
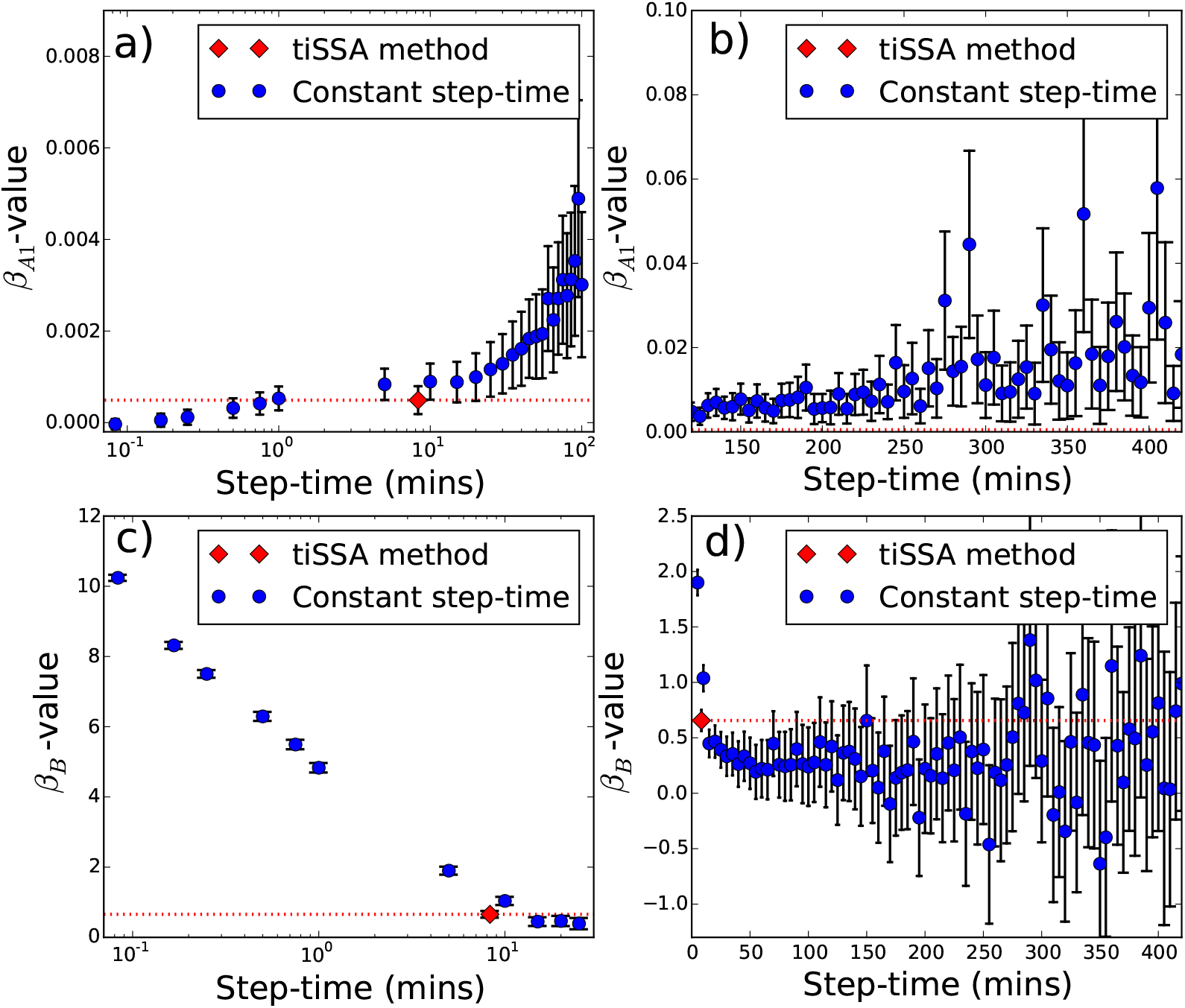
Results of step selection analysis on an example path, Goat Path 1. Panels (a) and (b) relate to the *β*_*A*1_ value, whereas Panels (c)d and (d) refer to *β*_B_. Recall that *β*_*A*1_ corresponds to the goat’s tendency to move toward the central point and *β*_B_ corresponds to the goat’s directional persistence. The blue dots show results from using iSSA with paths subsampled at regular step-times, whereas the red diamonds show results from analysing the paths using tiSSA. The horizontal location of the latter corresponds to the average step-time. Vertical bars represent 95% confidence intervals.

Across nearly all regularly subsampled paths we see that the inferred *β_B_*-values (Fig. 3b), representing the degree of directional persistence, are either inaccurate when compared to the inferred values from tiSSA (for small step-times) or so imprecise that the inferred *β_B_*-value is insignificant (for large step-times). For small constant step-times (those to the left of the red dot), *β_B_*-values increase rapidly with the subsampling frequency. However, all this means is that goats move in straight lines over very short time-periods, and as such it does not represent anything biologically meaningful about the tendency to choose short turns over longer ones.

When testing the time-dependent covariates (*Z*_*A*2_, *Z_D_* and *Z_E_*), the strength of directional persistence and tendency to orient turns towards the central point are both significantly higher than zero for all paths (Figure 4a,c). Figure 4b shows that the *β*_*A*2_-value, representing the tendency for the goats to move toward the central point with longer step-times, is significantly different from zero for four of the seven paths. All goats tend to have shorter step-times when the temperature is higher (Figure 4d). From field observations, we conjecture this is caused by the goats restricting their movements within shaded areas near the goat pen and other nearby buildings. Finally, all goat paths showed a tendency to have shorter step-times when moving through the rocky terrain (Fig. 4e). This is likely to be because it is more difficult to cross rocky terrain in a straight line than when crossing grassland or woodland, so the animals make more turns.

**Figure 4.**
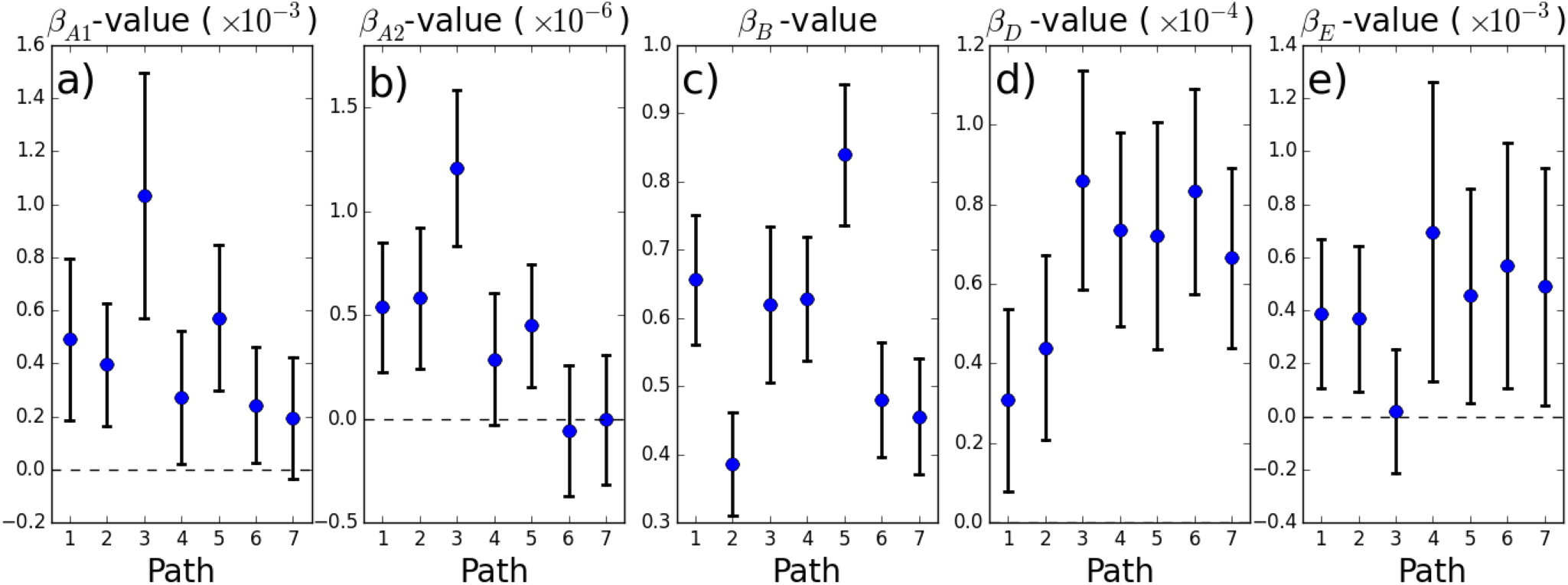
Coefficient values for (a) *β*_*A*1_, (b) *β*_*A*2_, (c) *β_B_*, (d) *β_D_*, and (e) *β_E_* with 95% confidence intervals, where *β*_*A*1_ represents the tendency of the goats to move toward the central point with greater strength the further they are from the central point, *β*_*A*2_ also represents the tendency to move toward the central point, with greater strength when further away from the central point and also greater step-times, *β_B_* represents the tendency to move in the same general direction, *β_D_* represents the tendency for the goats to have shorter step-times when the temperature is higher and *β_E_* represents the tendency for the goats to have shorter step-times when moving in rocky terrain.

## 4 Discussion

We have presented a method for step selection analysis with high resolution data, to help identify what drives animals to change their direction of movement. The method first finds the points at which an animal has made a turn, using the Turning-Points Algorithm (TPA) from Potts et al. (2018), then uses these points as locations in a modified version of iSSA, called time-varying iSSA (tiSSA). This generalises iSSA to allow for locations separated by differing time intervals, as arise in the output of TPA. Using both simulated paths and high frequency empirical data, we compare the tiSSA method to the usual use of iSSA, whereby decisions are inferred based on successive locations in low frequency data, showing that the latter can lead to both inaccurate and imprecise inference. Regularly subsampled paths at low frequencies tend to give both imprecise and inaccurate results whereas those at high frequencies tend to give precise but inaccurate results. This general trend follows for both the simulated and empirical datasets we used.

Being a direct generalisation of iSSA, and therefore SSA, our method can, in principle, be used to examine any of the behavioural processes affecting movement that have been demonstrated in the existing literature on step selection. These include attraction to areas of higher quality food (Merkle et al., 2016), avoidance of or attraction to linear features (Dickie et al., 2019), avoidance of prey (Latombe et al., 2014) or competitors (Vanak et al., 2013), territorial interactions (Potts et al., 2014a), effects of anthropogenic disturbances (Ladle et al., 2019), memory processes (Merkle et al., 2014), and much more (Thurfjell et al., 2014). However, tiSSA applied to high resolution data comes with the particular advantage that it is inferring movements that happen at points in space and time where the animal is known to have changed direction. Such changes cost energy (Wilson et al., 2013b) and so are likely to indicate distinct decisions by the animal (Potts et al., 2018). Therefore, the inference from tiSSA is likely to be more behaviourally meaningful than using SSA or iSSA on locations sampled at regular time-intervals.

The output of tiSSA also has the advantage of parametrising a model of animal movement, given by the movement kernel (Equation 1). This is also a feature of iSSA (and some predecessors, e.g. Potts et al. (2014b)), and is increasingly important for various applications, including finding mechanistic underpinnings of utilisation distributions (Signer et al., 2017), predicting aggregation and segregation phenomena (Potts and Schlägel, 2020), demarcating spatial scales of habitat selection (Bastille-Rousseau et al., 2015), predicting disease transmission rates (Merkle et al., 2018), and determining the energetic benefits of foraging strategies (Merkle et al., 2017). Our new technique thus opens up the possibility for such application based on the sort of high resolution data that is becoming increasingly prevalent in movement ecology (Williams et al., 2020).

As well as enabling such data to be used to answer questions already examined using iSSA with lower-resolution data, our tiSSA technique can also uncover processes that can cause animals to take turns more or less frequently. We have shown here that the goats in our study turn more frequently when the temperature is higher and the ground is rocky. In many behavioural changepoint studies, changes in turning frequency are viewed as changes in behavioural state (Patterson et al., 2008; Edelhoff et al., 2016). Whilst this may well be true, we have demonstrated that environmental conditions can also have an effect on turning frequency. This suggests it is important to account for such environmental effects in studies that seek to infer behavioural state from animal movement paths.

We have applied the TPA-tiSSA combination to data on dead-reckoned accelero-magnetometer data. However, the TPA-tiSSA technique is also applicable to any data that is of sufficiently high frequency for TPA to give an accurate reflection of the places that the animal actually turns. These data requirements are discussed in detail in Potts et al. (2018). In short, though, the key requirement is that there are sufficiently many recorded locations between each turning point for the sliding window in TPA to capture each turn. In practice, this will require at least a few dozen locations between successive turns with the magnetometers used here. However, if the noise in the data is lower, it may be possible to infer turning points with lower-frequency data. On the other hand, if noise is high, so that it is not possible to be confident about the precise locations of each turning-point (especially with respect to the value of the environmental covariates at each point), it may be advisable not to use tiSSA directly. One possible avenue would be to combine tiSSA with some sort of state-space modelling to account for the locational uncertainty, but we suspect that this is unlikely to be a trivial methodological extension. Also, it is entirely possible to use an alternative algorithm for calculating turning points instead of TPA, e.g. Codling and Plank (2011), if that turns out to be desirable.

Alongside application to high frequency data, tiSSA can in principle be used for any data where the steps are unevenly spaced in time. For example, many passerines tend to travel from tree to tree whilst foraging (Ellison et al., 2020). If one records the times and locations of a bird each time it switches tree then this would result in a sequence of locations, unevenly spaced in time, but where each ‘step’ from one location to the next represents a distinct decision by the animal. It would make sense to use tiSSA to understand the features of the trees that correlate to the decisions to leave one and move to another.

Many data sets also have measurements irregularly spaced in time due to limitations of tagging technology. However, unless there is reason to believe that the locations are gathered at behaviourally-meaningful points in time, we would recommend using continuous-time techniques instead of tiSSA (Wang et al., 2019), as continuous-time methods are capable of incorporating situations where decisions are made away from the measured locations (Blackwell et al., 2016). If the locations are gathered at behaviourally meaningful points in time, though, these continuous-time methods do not assume that these points are behaviourally significant, so do not make best use of the data. Another disadvantage of such techniques is that inference can often be prohibitively slow. However, there is evidence that this limitation is being overcome, thanks to improved statistical techniques, such as template model builder (Auger-Méthé et al., 2017; Jonsen et al., 2019).

In summary, our study provides the requisite methodological advance for meaningful use of step selection analysis with high frequency data, such as is becoming increasingly prevalent due to technological advances in both tagging hardware and software for post processing (Williams et al., 2020). Our study also demonstrates why this technique is advantageous in terms of producing accurate and precise results compared to using regularly subsampled, lower frequency data. As such, it opens the way for better inference for uncovering what drives animals to make movement decisions.

## Acknowledgments

RM acknowledges funding from a Leverhulme Trust Studentship as part of the Leverhulme Centre for Advanced Biological Modelling. Data collection was partly supported by a grant from the College of Science, Swansea University, for the ALPEN project to LB and RB, as well as Start-Up funding for LB (College of Science, Swansea University). We thank the goat herder, Gérard Perrier, and the Office Français de la Biodiversité (OFB; formerly, ONCFS) for logistical help during fieldwork. We thank Ulrike Schlägel, an anonymous reviewer, and the associate editor for very helpful comments on previous versions of the manuscript.

## Author Contributions

JRP, LB, and RPW conceived and designed the research; RM and JRP performed the research; LB, RPW, JR, RB and MG provided data; RM and JRP led the writing of the manuscript; all authors contributed critically to the drafts and gave final approval for publication.

## Data Accessibility

Data used in this manuscript will be archived on FigShare on acceptance of the manuscript.

