## Supplementary for "Why did the animal turn? Time-varying step selection analysis for inference between observed turning points in high frequency data"

#### S1 Parametrising the movement kernel using conditional logistic regression

Supposing that the location of an animal is at location $\mathbf{z}_{i-1}$, having arrived there on a heading of $\alpha_{\mathbf{z}_{i-1}}$ (i.e. the heading *prior* to being at location $\mathbf{z}_{i-1}$ is $\alpha_{\mathbf{z}_{i-1}}$). Then, by Equation (1) in the Main Text, the probability density of the animal making its next turn at location $\mathbf{x}$ and after time $\tau$ has elapsed is

$$f(\mathbf{x},\tau|\mathbf{z}_{i-1},\alpha_{\mathbf{z}_{i-1}}) = C^{-1}\phi(\mathbf{x},\tau|\mathbf{z}_{i-1})W(\mathbf{x},\tau,\mathbf{z}_{i-1},\alpha_{\mathbf{z}_{i-1}}) \quad (1)$$

where

$$C = \int_\Omega \int_0^\infty \phi(\mathbf{x},\tau|\mathbf{z}_{i-1})W(\mathbf{x},\tau,\mathbf{z}_{i-1},\alpha_{\mathbf{z}_{i-1}})\mathrm{d}\tau\mathrm{d}\mathbf{x} \quad (2)$$

is a normalising constant, which ensures that $f$ integrates to 1 over the domain of spatial availability, $\Omega$ and all possible step-times, $\tau$. Here, $W$ is in the form of Equation (4) in the Main Text, so depends on a vector of parameters, $\mathbf{B} = [\beta_1,\ldots,\beta_{n+4}]$. We aim to show that it is possible to parametrise $f$, and thus find $\mathbf{B}$, using conditional logistic regression (CLR).

The CLR technique involves first sampling $M$ times from $\phi(\mathbf{x},\tau|\mathbf{z}_{i-1})$. By Equation (2) from the Main Text, we can write $\phi$ as the product of three probability density functions $h(\alpha)$, $g_1(\tau)$ and $g_2(l|\tau)$, where $\alpha$ is the heading from $\mathbf{z}_{i-1}$ to $\mathbf{x}$ and $l$ is the distance between $\mathbf{z}_{i-1}$ and $\mathbf{x}$. So, to sample from $\phi$, requires us to first sample from $h(\alpha)$ and $g_1(\tau)$, which gives a set of paired control headings and step-times, $\{(\alpha'_{i,1}, \tau'_{i,1}), \dots, (\alpha'_{i,M}, \tau'_{i,M})\}$. Next, for each $\tau'_{i,j}$, we sample from $g_2(l|\tau'_{i,j})$ to give a step-length, $l'_{i,j}$ and define $\mathbf{x}'_{i,j} = \mathbf{z}_{i-1} + l'_{i,j}[\cos(\alpha'_{i,j}), \sin(\alpha'_{i,j})]$. Then the set of locations at the ends of control steps and their corresponding step-times is $\{(\mathbf{x}'_{i,1}, \tau'_{i,1}), \dots, (\mathbf{x}'_{i,M}, \tau'_{i,M})\}$.

Suppose the path of an animal is given by the set of turning points (locations) and headings $\mathbf{S} = \{(\mathbf{z}_0, \dots, \mathbf{z}_N), (\tau_1, \dots, \tau_N)\}$, where $\mathbf{z}_i$ is the $i^{\text{th}}$ turning-point and $\tau_i$ is the step-time between $\mathbf{z}_{i-1}$ and $\mathbf{z}_i$. Then the likelihood of $\mathbf{B}$ given $\mathbf{S}$ for CLR is

$$L_{CLR}(\mathbf{B}|\mathbf{S}) = \prod_{i=2}^{N} \frac{W(\mathbf{z}_i, \tau_i, \mathbf{z}_{i-1}, \alpha_{\mathbf{z}_{i-1}})}{\sum_{j=0}^{M} W(\mathbf{x}'_{i,j}, \tau'_{i,j}, \mathbf{z}_{i-1}, \alpha_{\mathbf{z}_{i-1}})} \tag{3}$$

where $\alpha_{\mathbf{z}_{i-1}}$ is the heading from $\mathbf{z}_{i-2}$ to $\mathbf{z}_{i-1}$ (note that this is why the indexing of $i$ starts at two) and we define $\mathbf{x}'_{i,0} = \mathbf{z}_i$ and $\tau'_{i,0} = \tau_i$.

Showing that CLR can be used to parametrise Equation (1) requires us to show that Equation (3) is maximised at the same value of $\mathbf{B}$ as the following likelihood function

$$\begin{aligned} L_I(\mathbf{B}|\mathbf{S}) &= \prod_{i=2}^{N} f(\mathbf{z}_i, \tau_i | \mathbf{z}_{i-1}, \alpha_{\mathbf{z}_{i-1}}) \\ &= \prod_{i=2}^{N} \frac{\phi(\mathbf{z}_i, \tau_i|\mathbf{z}_{i-1}) W(\mathbf{z}_i, \tau_i, \mathbf{z}_{i-1}, \alpha_{\mathbf{z}_{i-1}})}{\int_\Omega \int_0^\infty \phi(\mathbf{x}, \tau|\mathbf{z}_{i-1}) W(\mathbf{x}, \tau, \mathbf{z}_{i-1}, \alpha_{\mathbf{z}_{i-1}}) \, \mathrm{d}\tau \, \mathrm{d}\mathbf{x}}. \end{aligned} \tag{4}$$

The central observation is that the denominator in Equation (4) can be approximated, up to a multiple that is constant and with respect to $\beta_1, \dots, \beta_{n+4}$, by the denominator of Equation (3). This follows from the theory of Monte-Carlo integration. In addition, since $\phi(\mathbf{z}_i, \tau_i|\mathbf{z}_{i-1})$ does not depend on $\beta_1, \dots \beta_{n+4}$, we can therefore write

$$\frac{\phi(\mathbf{z}_i, \tau_i|\mathbf{z}_{i-1}) W(\mathbf{z}_i, \tau_i, \mathbf{z}_{i-1}, \alpha_{\mathbf{z}_{i-1}})}{\int_\Omega \int_0^\infty \phi(\mathbf{x}, \tau|\mathbf{z}_{i-1}) W(\mathbf{x}, \tau, \mathbf{z}_{i-1}, \alpha_{\mathbf{z}_{i-1}}) \, \mathrm{d}\tau \, \mathrm{d}\mathbf{x}} \approx K_i \frac{W(\mathbf{z}_i, \tau_i, \mathbf{z}_{i-1}, \alpha_{\mathbf{z}_{i-1}})}{\sum_{j=0}^{M} W(\mathbf{x}'_{i,j}, \tau'_{i,j}, \mathbf{z}_{i-1}, \alpha_{\mathbf{z}_{i-1}})} \tag{5}$$

where $K_i$ is constant with respect to $\beta_1, \dots, \beta_{n+4}$. Equation (5) shows that the $\beta_i$-values that maximise Equation (4) are approximately equal to those $\beta_i$-values that maximise Equation (3). The approximation becomes exact as $M \to \infty$. Consequently, we can use CLR to find the $\mathbf{B}$ that maximises $L_I(\mathbf{B}|\mathbf{S})$ and hence find the maximum likelihood approximation for parameters, $\mathbf{B}$, of the model in Equation (1).

#### S2 Fitting the $k_2$ and $\theta_2$ parameters to $g_2^{(r)}(l|\tau)$

The conditional logistic regression method requires step-lengths to be sampled from the probability density function, $g_2(l|\tau)$ (Section S1). By Equation (5) in the Main Text we draw step-lengths from the following probability density function

$$g_2^{(r)}(l|\tau) = \frac{l^{k_2-1}\exp(-l/(\theta_2\tau))}{\Gamma(k_2)(\theta_2\tau)^{k_2}}. \tag{6}$$

Since the right-hand-side of (6) is a different gamma distribution for each $\tau$, it is not straightforward to fit (6) to data using out-of-the-box code (e.g. `fitdistr` in R).

However, we can instead fit a gamma distribution to the step-speeds to parametrise Equation (6). To explain why, first note that the likelihood function for the model in Equation (6) is

$$\begin{aligned} L_l(k_2, \theta_2|l_1, \ldots, l_N, \tau_1, \ldots, \tau_N) &= \prod_{i=1}^{N} \frac{l_i^{k_2-1}\exp(-l_i/(\theta_2\tau_i))}{\Gamma(k_2)(\theta_2\tau_i)^{k_2}} \\ &= \left(\prod_{j=1}^{N} \frac{1}{\tau_j}\right) \prod_{i=1}^{N} \frac{s_i^{k_2-1}\exp(-s_i/\theta_2))}{\Gamma(k_2)\theta_2^{k_2}}. \end{aligned} \tag{7}$$

where $s_i = l_i/\tau_i$ is the step-speed.

Now, the likelihood function for fitting a gamma distribution to the step-speeds is

$$L_s(k_2, \theta_2|s_1, \ldots, s_N) = \prod_{i=1}^{N} \frac{s_i^{k_2-1}\exp(-s_i/\theta_2))}{\Gamma(k_2)\theta_2^{k_2}}. \tag{8}$$

Notice that $L_l$ and $L_s$ are each maximised by the same values of $k_2$ and $\theta_2$. Therefore we can parametrise Equation (6) by fitting a single gamma distribution to $s_1, \ldots, s_N$ (e.g. `fitdistr` in R), then setting $k_2$ and $\theta_2$ to be the shape and scale values of the best-fit gamma distribution of step-speeds.

### S3 Central area identification

In the Main Text, we include a covariate ($Z_{A1}$) which we use to test the hypothesis that the goats have a tendency to move toward a central area. This hypothesis is represented by an attraction to a single point, $\mathbf{x}_{cp}$, at the centre of the area. We identify this point for each goat as the centre of the site of interest found using the Sites of Interest Algorithm from Munden et al. (2019). Here, we summarise the key concepts from this algorithm and explain how we applied it to our particular case.

The Sites of Interest Algorithm works by first sliding a circle of fixed radius, $R$, along an animal's path. For each location the circle is centred at, the amount of time the animal spends inside the circle is calculated, which we term the *usage time*. Next, the set of circles is rarefied by removing any that overlap with a circle of a higher usage time. We then rank the remaining circles in terms of usage time and calculate the percentage drop in usage time between successive circles in this ranked list. The point at which this percent drop is maximised (the *maximum percent drop*) is used to demarcate "sites" (those prior to the maximum percent drop) and "non-sites" (those after the maximum percent drop). This method is analogous to using a scree plot in principal component analysis: see Munden et al. (2019) for details.

The question the method raises is how to choose $R$. For all the values of $R$ (we used $10, 15, \ldots, 100$m), across all paths, the Sites of Interest Algorithm found either a single site encompassing the goat pen and salt licks (72% of cases; Fig. SF1) or a cluster of sites all close to the same central area. To define the Site of Interest, we used the value of $R$ that gave the highest maximum percent drop out of those tested.

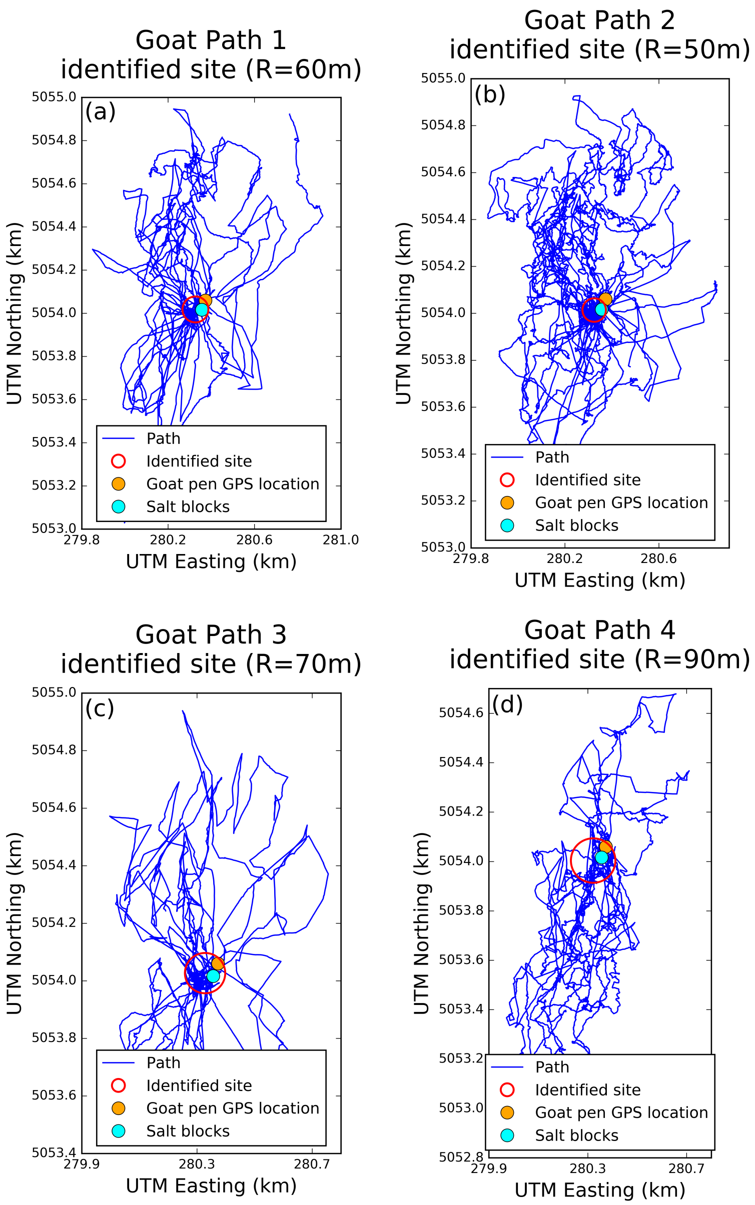


**Figure SF1.** Some example trajectories of individual goats with the single identified site of interest and the GPS location of the goat pen.

#### S4 Step-time covariate models

Here we write out the functional forms for the movement kernel under the Straight Returns Model ($f_2^{(r)}$), the Temperature Model ($f_3^{(r)}$) and the Rocky Terrain Model ($f_4^{(r)}$) as

$$f_2^{(r)}(\mathbf{x},\tau|\mathbf{z},\alpha_{\mathbf{z}}) \propto \frac{1}{\tau}\exp\Bigg(\tilde{\beta}_1^{(r)}\tilde{Z}_1(\mathbf{x},\mathbf{z},\tau) + \beta_2^{(r)} Z_2(\mathbf{x},\mathbf{z},\alpha_{\mathbf{z}}) + \beta_3^{(r)} Z_3(\mathbf{x},\mathbf{z}) + \left(\beta_6^{(r)} - \frac{1}{\theta_1}\right)\tau + \left(\beta_7^{(r)} - \frac{1}{\theta_2}\right)\frac{|\mathbf{x}-\mathbf{z}|}{\tau} + (\beta_8^{(r)} + k_1 - 1)\ln\tau + (\beta_9^{(r)} + k_2 - 1)\ln\left(\frac{|\mathbf{x}-\mathbf{z}|}{\tau}\right)\Bigg) \tag{9}$$

$$f_3^{(r)}(\mathbf{x},\tau|\mathbf{z},\alpha_{\mathbf{z}}) \propto \frac{1}{\tau}\exp\Bigg(\beta_1^{(r)} Z_1(\mathbf{x},\mathbf{z}) + \beta_2^{(r)} Z_2(\mathbf{x},\mathbf{z},\alpha_{\mathbf{z}}) + \beta_3^{(r)} Z_3(\mathbf{x},\mathbf{z}) + \beta_4^{(r)} Z_4(\mathbf{x},\mathbf{z},\tau) + \left(\beta_6^{(r)} - \frac{1}{\theta_1}\right)\tau + \left(\beta_7^{(r)} - \frac{1}{\theta_2}\right)\frac{|\mathbf{x}-\mathbf{z}|}{\tau} + (\beta_8^{(r)} + k_1 - 1)\ln\tau + (\beta_9^{(r)} + k_2 - 1)\ln\left(\frac{|\mathbf{x}-\mathbf{z}|}{\tau}\right)\Bigg) \tag{10}$$

$$f_4^{(r)}(\mathbf{x},\tau|\mathbf{z},\alpha_{\mathbf{z}}) \propto \frac{1}{\tau}\exp\Bigg(\beta_1^{(r)} Z_1(\mathbf{x},\mathbf{z}) + \beta_2^{(r)} Z_2(\mathbf{x},\mathbf{z},\alpha_{\mathbf{z}}) + \beta_3^{(r)} Z_3(\mathbf{x},\mathbf{z}) + \beta_5^{(r)} Z_5(\mathbf{x},\tau) + \left(\beta_6^{(r)} - \frac{1}{\theta_1}\right)\tau + \left(\beta_7^{(r)} - \frac{1}{\theta_2}\right)\frac{|\mathbf{x}-\mathbf{z}|}{\tau} + (\beta_8^{(r)} + k_1 - 1)\ln\tau + (\beta_9^{(r)} + k_2 - 1)\ln\left(\frac{|\mathbf{x}-\mathbf{z}|}{\tau}\right)\Bigg) \tag{11}$$

### S5 Additional Tables

| | Step-Time | | Step-Speed | |
|---|---|---|---|---|
| Path | Shape ($k_1$) | Scale ($\theta_1$) | Shape ($k_2$) | Scale ($\theta_2$) |
| 1 | 0.50 | 1007 | 0.37 | 0.26 |
| 2 | 0.43 | 804 | 0.48 | 0.26 |
| 3 | 0.63 | 933 | 0.57 | 0.12 |
| 4 | 0.34 | 1138 | 0.31 | 0.32 |
| 5 | 0.43 | 1045 | 0.47 | 0.20 |
| 6 | 0.39 | 927 | 0.44 | 0.25 |
| 7 | 0.33 | 1092 | 0.38 | 0.32 |

**Table ST1.** Fitted shape and scale values for a gamma distribution to the step times and step speeds inferred from the paths rarefied using the Turning Points Algorithm.

| True $\lambda$ | True $\beta$ | Inferred $\beta$ with correction | Inferred $\beta$ without correction |
|---|---|---|---|
| 0.1 | 0.5 | (0.48,0.56) | (0.51,0.58) |
| 0.1 | 1 | (0.97, 1.05) | (1.03, 1.11) |
| 0.1 | 2 | (1.90,2.10) | (2.06,2.15) |
| 0.2 | 0.5 | (0.43,0.58) | (0.44,0.58) |
| 0.2 | 1 | (1.00,1.16) | (1.09,1.24) |
| 0.2 | 2 | (1.91,2.14) | (2.19,2.38) |
| 0.2 | 0.5 | (0.43,0.60) | (0.53,0.68) |
| 0.5 | 1 | (1.01,1.20) | (1.11,1.27) |
| 0.5 | 2 | (1.82,2.10) | (2.01,2.18) |

**Table ST2.** Results of inferring $\beta$ from the simulated paths using tiSSA with a correction for the step-length and step-time (as advised), compared with neglecting this correction. The first two columns give the true values of $\lambda$ and $\beta$ respectively. The third (resp. fourth) column gives the 95% confidence intervals of using tiSSA with (resp. without) the correction terms.

### S6 Additional Figures

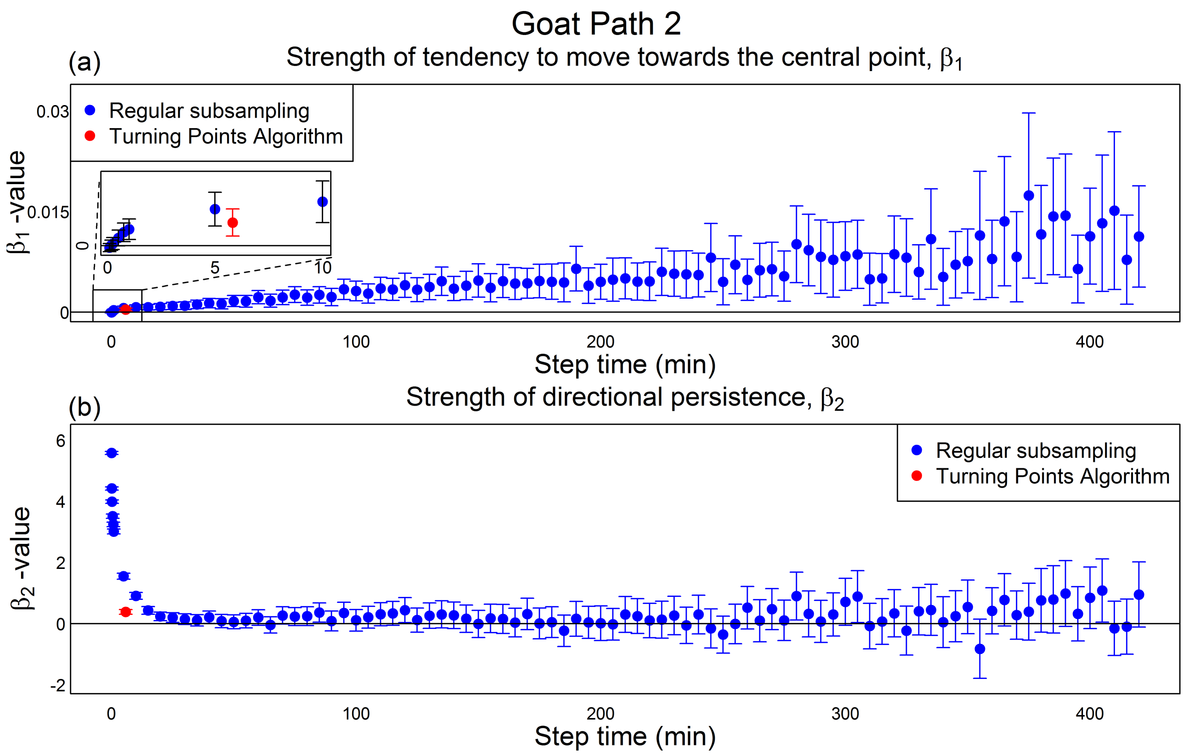


**Figure SF2.** Results of step selection analysis on an example path, Goat Path 2. Panels (a) and (b) relate to the $\beta_1$ and $\beta_2$ values, respectively. Recall that $\beta_1$ corresponds to the goat's tendency to move toward the central point and $\beta_2$ corresponds to the goat's directional persistence. Results from regular subsampling are presented in blue and the Turning-Points Algorithm results in red. The horizontal location of the latter corresponds to the average step-time. Vertical bars represent 95% confidence intervals.

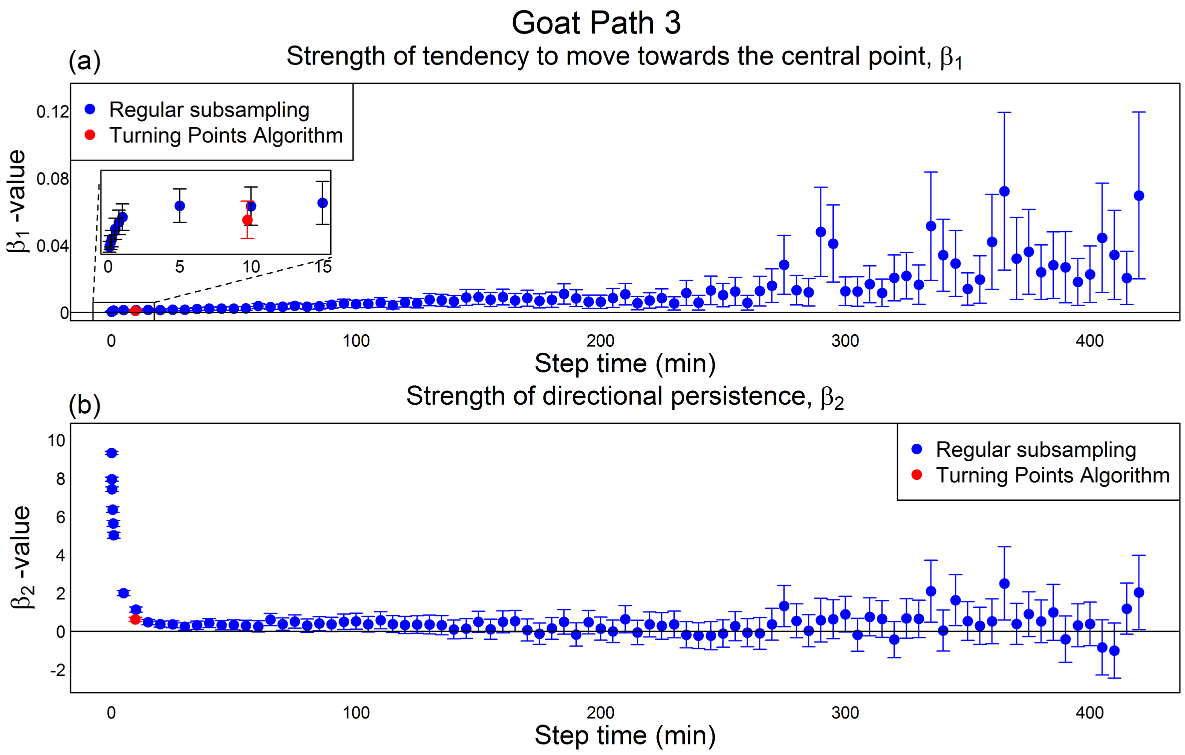


**Figure SF3.** Results of step selection analysis on an example path, Goat Path 3. Format is identical to Figure SF2.

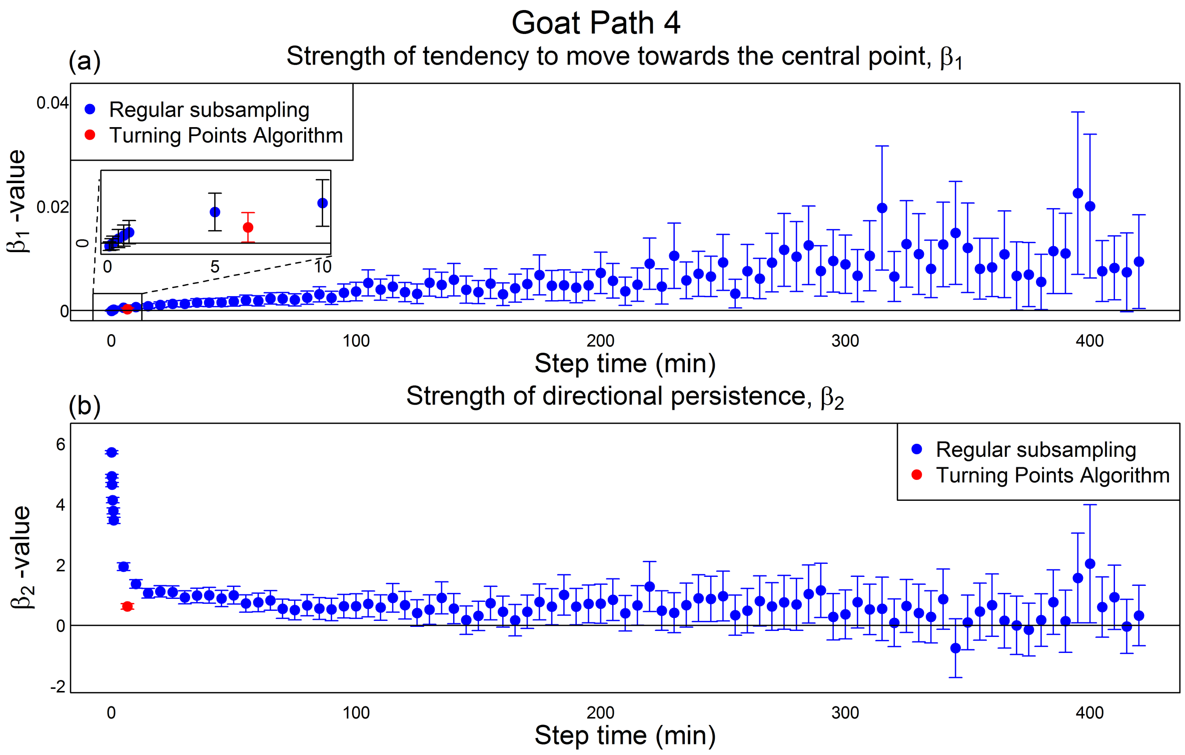


**Figure SF4.** Results of step selection analysis on an example path, Goat Path 4. Format is identical to Figure SF2.

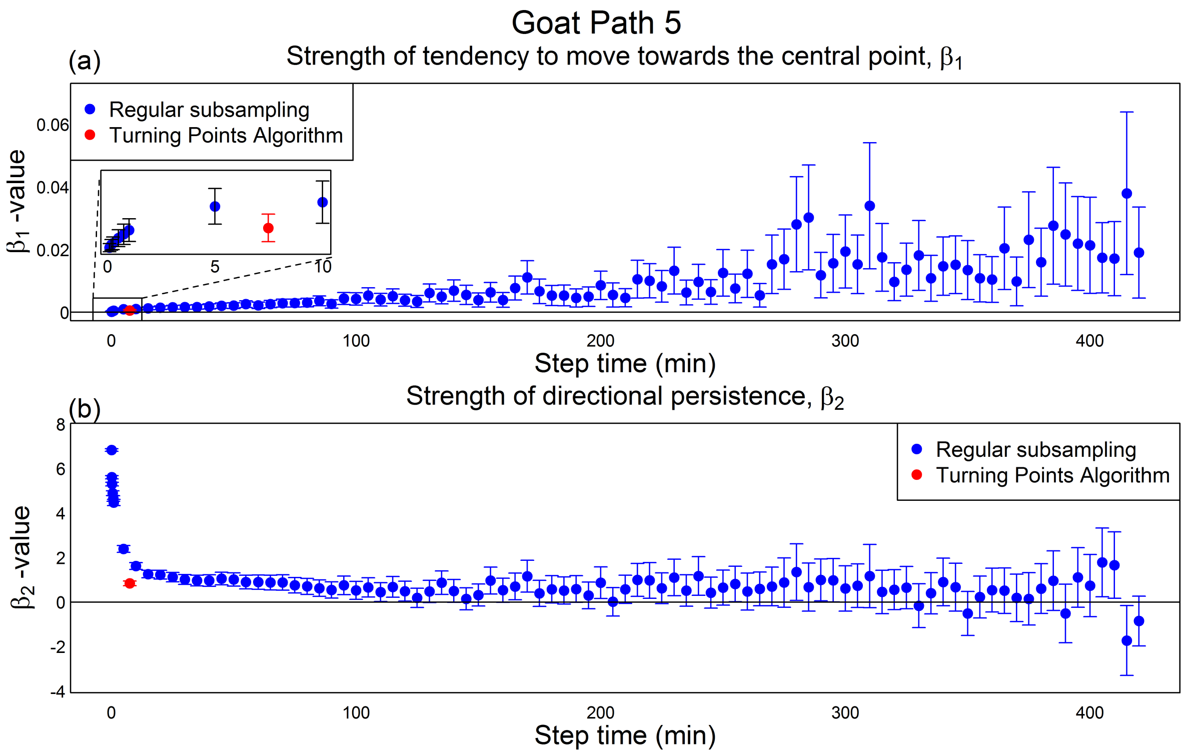


**Figure SF5.** Results of step selection analysis on an example path, Goat Path 5. Format is identical to Figure SF2.

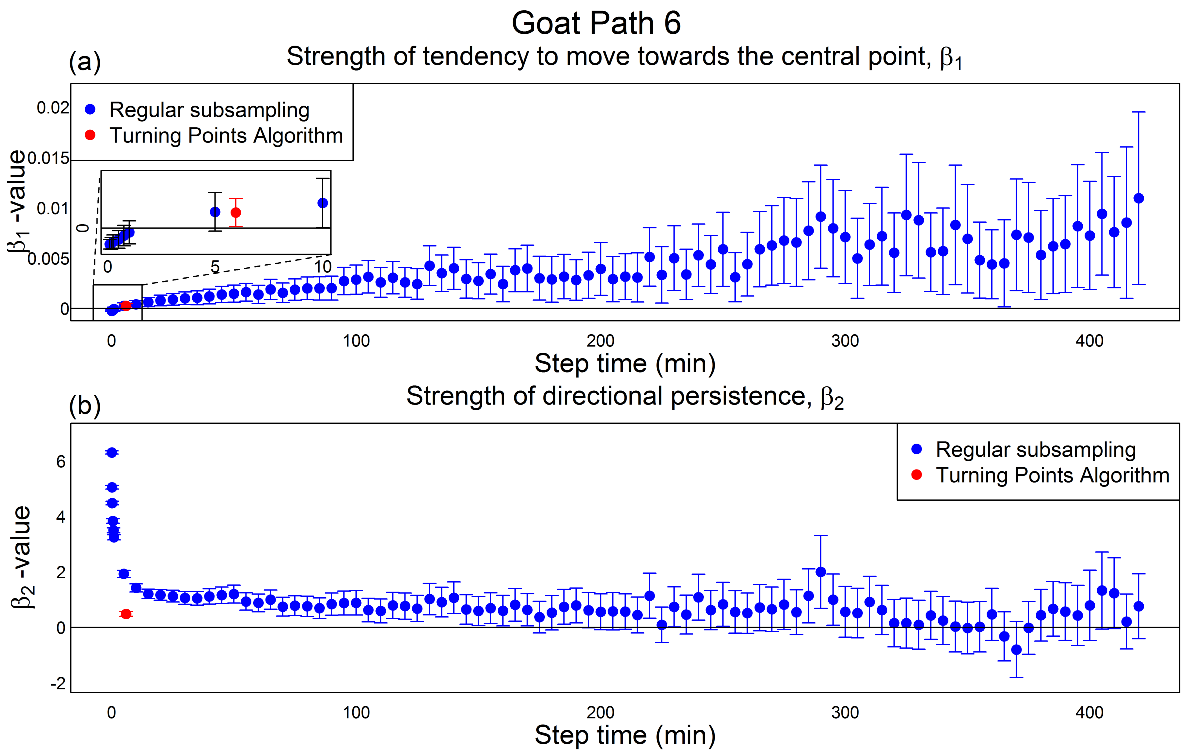


**Figure SF6.** Results of step selection analysis on an example path, Goat Path 6. Format is identical to Figure SF2.

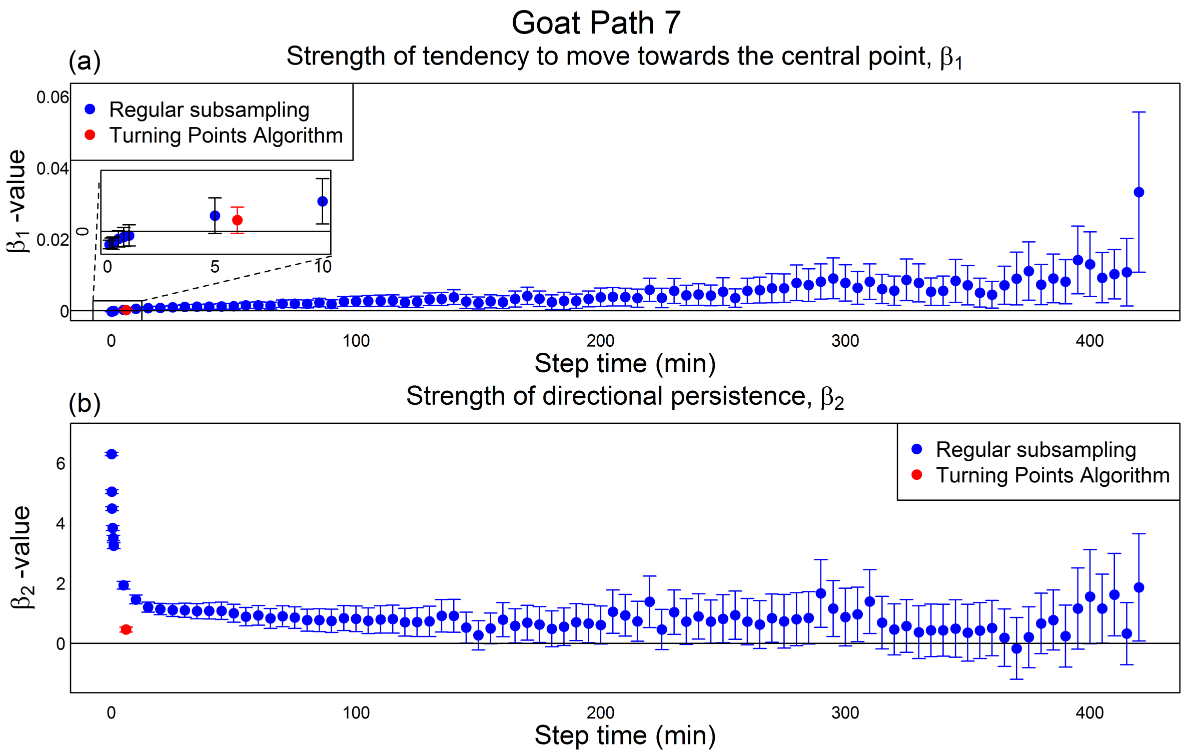


**Figure SF7.** Results of step selection analysis on an example path, Goat Path 7. Format is identical to Figure SF2.

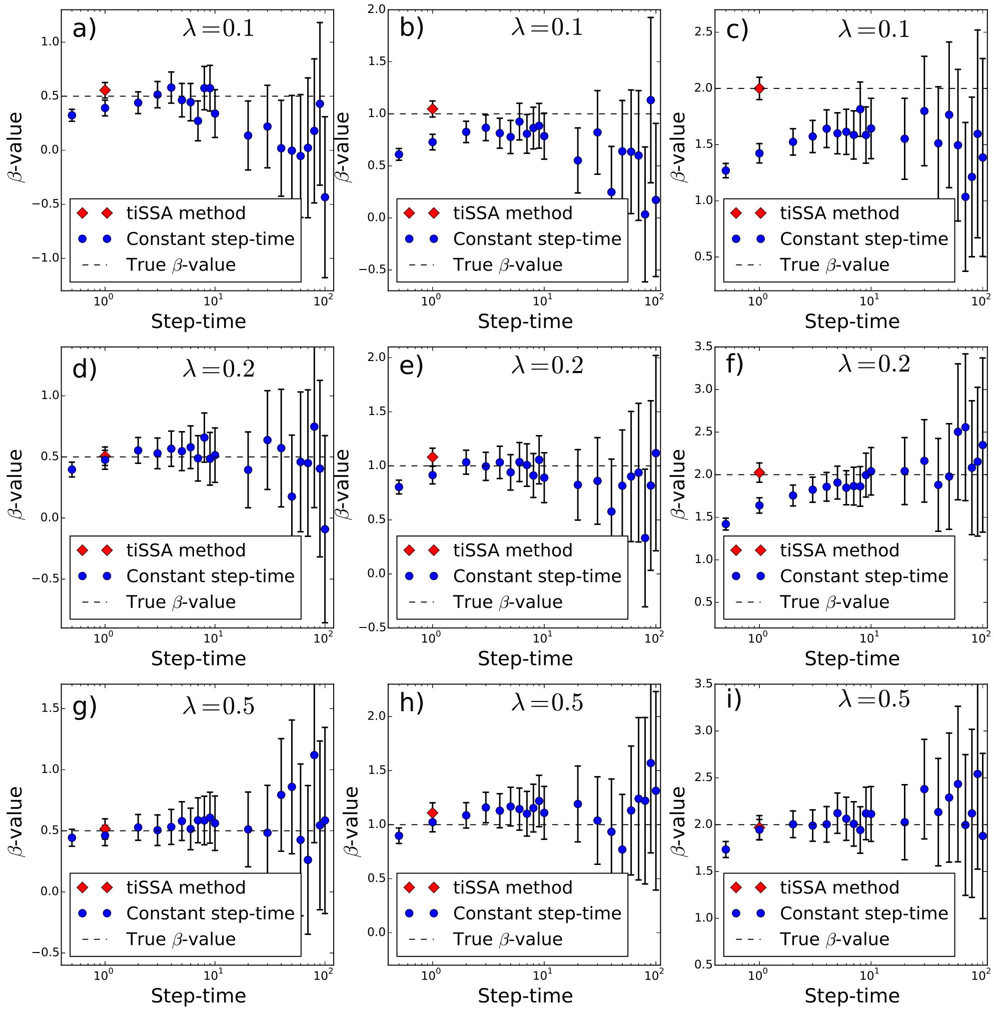


**Figure SF8.** Results of the simulation analysis. The panels in the left, middle, and right columns show, respectively, results where $\beta = 0.5$, $\beta = 1$, and $\beta = 2$. The panels in the top, middle, and bottom rows show, respectively, results where $\lambda = 0.1$, $\lambda = 0.2$, and $\lambda = 1$. The red diamonds show results from analysing the paths using tiSSA, whereas the blue dots show results from using iSSA with paths subsampled at regular step-times. Error bars show 95% confidence intervals.
